# Walking bumblebees see faster

**DOI:** 10.1101/2022.12.20.521190

**Authors:** Lisa Rother, Robin Müller, Erwin Kirschenmann, Sinan Kaya-Zeeb, Markus Thamm, Keram Pfeiffer

**Author notes:** shared first author (Lisa Rother, Robin Müller).

## Abstract

The behavioral state of an animal has profound effects on neuronal information processing. Locomotion changes the response properties of visual interneurons in the insect brain, but it is still unknown if it also alters the response properties of photoreceptors. Photoreceptor responses become faster at higher temperatures. It has therefore been suggested that thermoregulation in bees and other insects could improve temporal resolution in vision, but direct evidence for this idea has so far been missing. Here we compared electroretinograms (ERGs) from the compound eyes of tethered bumblebees that were either sitting still or were walking on an air supported ball. We found that the visual processing speed strongly increased when the bumblebees were walking. By monitoring the bumblebees’ eye temperature during our recordings, we saw that the increase in response speed was in synchrony with a rise of the eye temperature. By artificially heating the bee’s head, we show that the walking induced temperature increase of the visual system is sufficient to explain the rise in processing speed. We also show that walking accelerates the visual system to the equivalent of a 14-fold increase in light intensity. We conclude that the walking-induced rise in temperature accelerates the processing of visual information in bumblebees, which is an ideal strategy to process the increased information flow during locomotion.

## Introduction

It is well documented that the processing of visual information is adjusted to the behavioral state in humans (Cao and Händel 2019), rodents (Murakami et al. 2005; Cano et al. 2006; Niell and Stryker 2010; Maimon 2011; Polack et al. 2013; Christensen and Pillow 2022), and insects (Maimon 2011; Cheng and Frye 2020). Motion-sensitive neurons in flies increase their gain during walking (Chiappe et al. 2010), flying (Maimon et al. 2010) and movement of the halteres (Rosner et al. 2010) and these locomotion-dependent changes are mediated by octopamine (Rien et al. 2012; Suver et al. 2012). However, it remains unclear if the behavioral state effects only the neuronal processing, or if locomotion might also alter sensory processing at the photoreceptor level. Previous studies have shown that photoreceptors of different insect species respond faster at higher temperatures (Kikuchi et al. 1961; French and Järvilehto 1978; Weckström et al. 1985; Roebroek et al. 1990; Tatler et al. 2000). Bumblebees, like other insect species are capable of regulating their body temperature over a wide range of temperatures independent of the ambient temperature by producing heat using their flight muscles. The heat generated in the thorax is then passed to the head by anterograde pumping of hemolymph (Heinrich 1993). It has therefore been suggested that insects that are capable of thermoregulation, might be able to use this ability to increase the temporal resolution of their visual systems (Tatler et al. 2000). This would be similar to some species of large predatory pelagic fish that are able to keep their eye temperature 10-15°C above the ambient water temperature, which results in increased temporal resolution of their visual system (Block 1986, Fritsches et al. 2005). However, direct experimental evidence showing that locomotion increases the processing speed of the visual system in insects has been lacking so far. By comparing electroretinograms (ERGs) from tethered bumblebees that were either walking or sitting, we provide here the first evidence that locomotion indeed speeds up the visual processing in bumblebees.

## Results

### Walking accelerates the ERG response

We first measured the ERG in response to individual light pulses produced by a green (530 nm) LED (Figure 1B; 50 ms duration at 1 Hz, pulse intensity: 2*10^15^ photons/cm^2^*s) prior to walking (sitting, light blue), during walking (violet), and after walking (sitting, dark blue) (Fig. 1B). We found that the time course of the ERG was shifted to the left during walking in comparison to the ERGs measured while the animal was sitting prior to and after walking. To quantify this shift, we measured the time between the onset of the stimulus and the time at which the ERG response crossed a threshold value of −2.5 mV (Figure 1B). Across all recordings (n = 16), the median time to threshold was significantly shorter when the animals were walking (6.7 ms) than when they were sitting (before walking: 7.8 ms, after walking: 7.5 ms; Figure 1C; before walking vs. walking: p = 8*10^−4^, after walking vs walking; p = 4*10^−4^, before walking vs. after walking: p = 0.061, Wilcoxon-Test with Bonferroni correction (α = 0.0167)). This suggests a locomotor-dependent increase in visual response speed in walking bumblebees.

**Figure 1:**
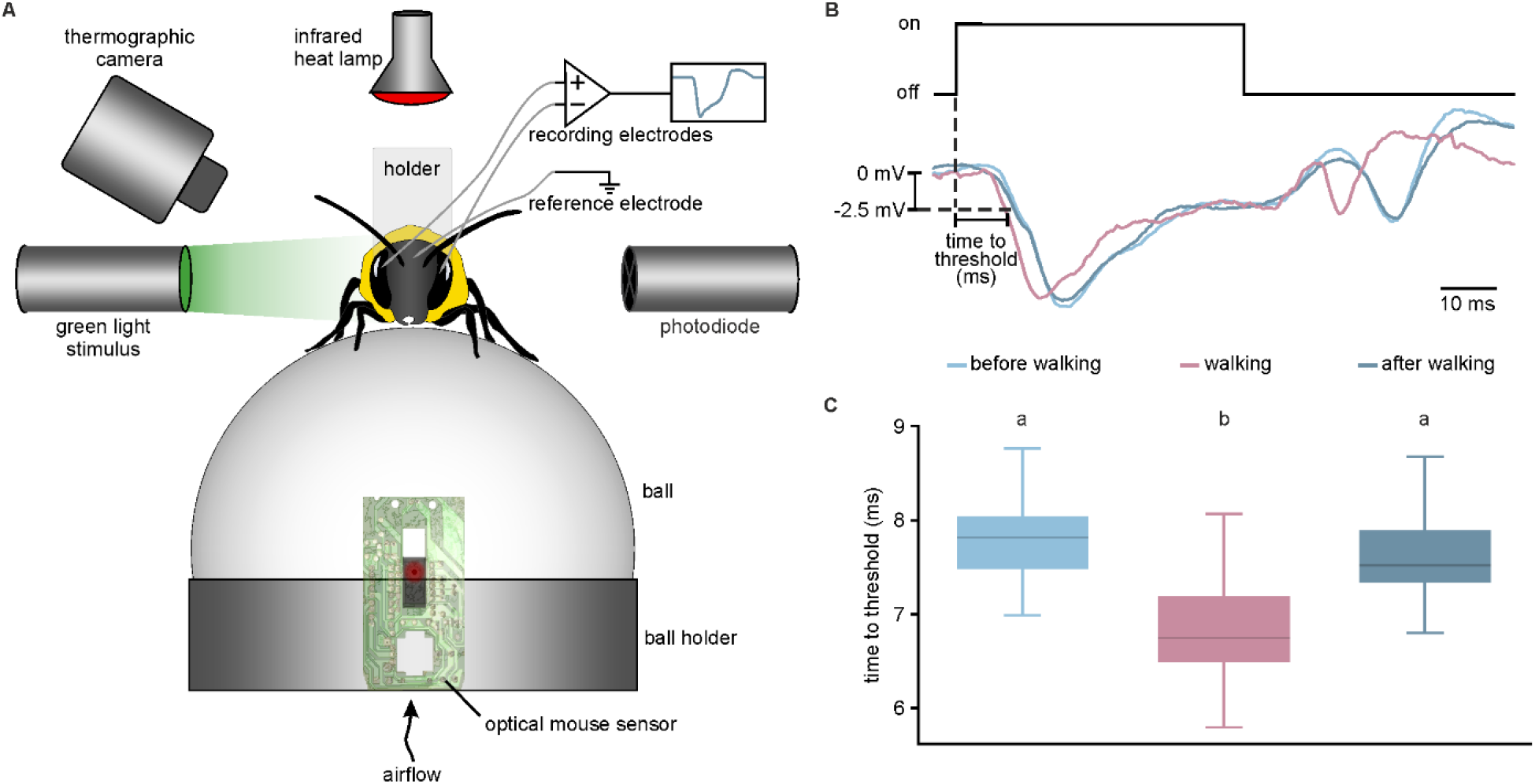
Experimental setup and electroretinograms (ERGs) of sitting and walking bumblebees stimulated with green light pulses. (A) Schematic illustration of the experimental setup. The bumblebee was tethered and positioned on top of an air-supported ball. While presenting green light (530 nm) pulses, the ERG was recorded. An infrared lamp and a thermographic camera were used to control and measure the temperature. (B) Upper trace: single pulse of green light (50 ms duration at 1 Hz, pulse intensity: 2*10^15^ photons/cm^2^*s). Lower panel: ERG response curves of one example trace of an animal during walking (violet) or sitting (light blue: before walking, dark blue: after walking). The time difference between the onset of the light pulse and the crossing of a threshold of −2.5 mV gave the time to threshold. (C) Boxplots of the time to threshold of the ERG response during walking (violet) or sitting (light blue: before walking, dark blue: after walking) of 16 bumblebees. The time to threshold was significantly shorter during walking than during sitting (p < 0.001, Wilcoxon-Test with Bonferroni correction), but no differences were seen when comparing before and after walking (p = 0.061, Wilcoxon-Test with Bonferroni correction). Boxplots show median, interquartile range (IQR), whiskers with 1.5x IQR and outliers >1.5x IQR.

### Increased visual response speed during walking coincides with increase in eye temperature

To be able to assess the frequency range of the ERG response that was particularly affected by walking, we repeated our experiments, this time using a Gaussian white noise stimulus (Figure 2A, Methods and Supplements for details). This allowed us to continuously monitor the cross-correlation between the stimulus signal and the ERG as well as to compute the frequency response function. The cross-correlation computes the similarity of two curves, and its minimum (or maximum) indicates the relative shift of the two curves, i.e. the time lag between them. Figure 2B displays example traces of cross-correlation curves showing that the time lag was shorter during walking (violet, 6.5 ms) than during sitting both before walking and after walking (light and dark blue, both 8.3 ms). This observation was confirmed in all tested animals, and the difference in time lag was significant (Figure 2C; n = 12; median values: before walking, 9.0 ms, walking, 7.4 ms, after walking, 9.0 ms; before walking vs. walking: p = 0.002, after walking vs. walking: p = 0.002, before walking vs. after walking: p = 0.45, Wilcoxon-Test with Bonferroni correction (α = 0.0167)).

**Figure 2:**
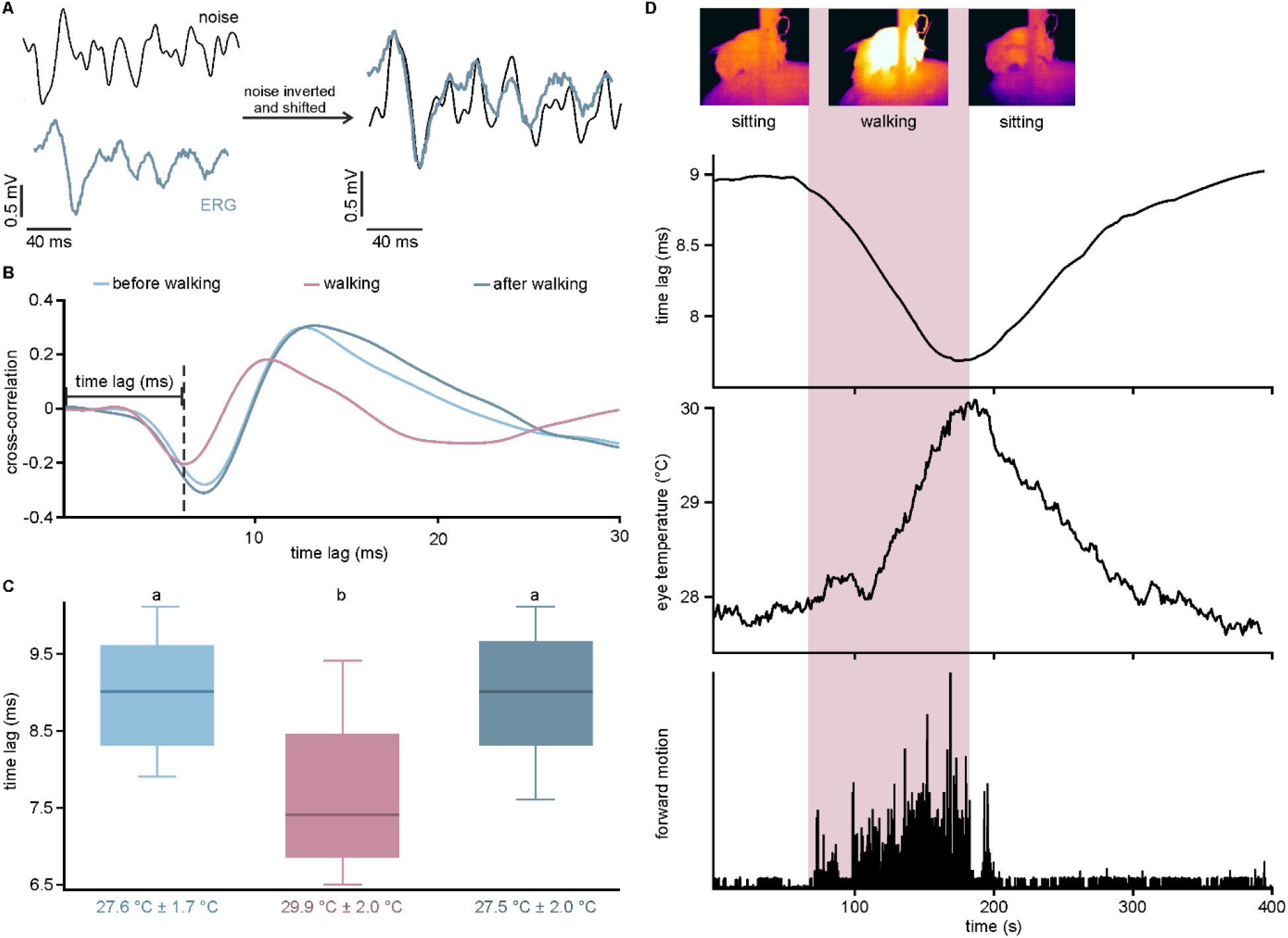
Effects of locomotion on the ERG of *B. terrestris* during stimulation with Gaussian white noise. (A) Section of the Gaussian noise stimulus (black; for whole trace, see supplementary information) and the corresponding ERG response (blue). To illustrate the similarity between noise stimulus and ERG response, the noise track was inverted and shifted (right). (B) Cross-correlation between the signal of the photodiode and the ERG to measure the time lag of the ERG during stimulation with Gaussian noise. Cross-correlation measures the similarity of two waveforms in the time domain and can assume values between −1 (identical waveforms with inverted signs) and 1 (identical waveforms). The x-value at the minimum of the cross-correlation indicates the time lag between the two curves. The minimum during walking (violet) occurs earlier than when the animal is sitting (blue). (C) Boxplots show the responses of 12 animals to the noise track, before walking (light blue; 27.6°C ± 1.7°C), during walking (violet; 29.9°C ± 2.0°C), and after walking (dark blue; 27.5°C ± 2.0°C). The time lag (ms) was significantly shorter when walking than sitting (p = 0.002, Wilcoxon-Test with Bonferroni correction), but no differences were seen when comparing before and after walking (p = 0.45, Wilcoxon-Test with Bonferroni correction). Boxplots show median, interquartile range (IQR), whiskers with 1.5x IQR and outliers >1.5x IQR. (D) Time course of time lag (upper trace), temperature (middle trace), and forward motion (lower trace). During locomotion (highlighted in violet), the time lag decreased and the temperature increased (symbolized with thermographic pictures at the top). After walking, the values return to their initial level.

To evaluate the frequency range of the ERG response that was specifically affected we calculated the gain and the linear coherence (Fig. 3A, n = 11) between the stimulus and the ERG. Gain is a measure of how strong an output signal (here: Gaussian white noise light stimulus) is compared to the input signal (here: ERG), at each frequency in the signal. Linear coherence is a value between 0 and 1 that measures the similarity in frequency content of two signals. It is 1, if the relationship between the input and the output (ERG) is completely linear and no noise is added by the system (i.e. the visual system). We found that both gain and coherence were on average shifted to higher frequencies during walking (Figure 3A). In some recordings such a shift was not observed and this was more likely in animals that heated up less during walking (Supplemental Figure 7, left). To quantify the frequency-dependent changes that occurred during walking, we fitted second order low-pass filter functions to the gain (see Methods for details). The time constant τ of this filter determines at which frequency the filter function rolls of such that smaller values of τ indicate a shift to higher roll-off frequencies. Comparing the median time constants during sitting and walking, we found, that τ became significantly smaller during walking (Figure 3B middle panel; n = 11; median values: before walking, 1.41 ms, walking, 1.25 ms, after walking, 1.39 ms; before walking vs. walking: p = 0.001, after walking vs. walking: p = 0.001, before walking vs. after walking: p = 0.014, Wilcoxon-Test with Bonferroni correction (α = 0.0167)). We also found a small, but significant difference in τ between the two sitting conditions, which probably indicates that the system had not completely recovered from the walking condition. Walking had no effect on the damping factor ζ, but as for τ, there was a small, but significant difference in the values of ζ between the two sitting conditions (Figure 3B lower panel; n = 11; median values: before walking, 0.150, walking, 0.146, after walking, 0.148; before walking vs. walking: p = 0.563, after walking vs. walking: p = 0.123, before walking vs. after walking: p = 0.004, Wilcoxon-Test with Bonferroni correction (α = 0.0167)). The amplitude β of the fit did not differ between any of the conditions (Figure 3B upper panel; n = 11; median of normalized values: before walking, 0.955, walking, 0.996, after walking, 0.939; before walking vs. walking: p = 0.700, after walking vs. walking: p = 0.064, before walking vs. after walking: p = 0.320, Wilcoxon-Test with Bonferroni correction (α = 0.0167)).

**Figure 3:**
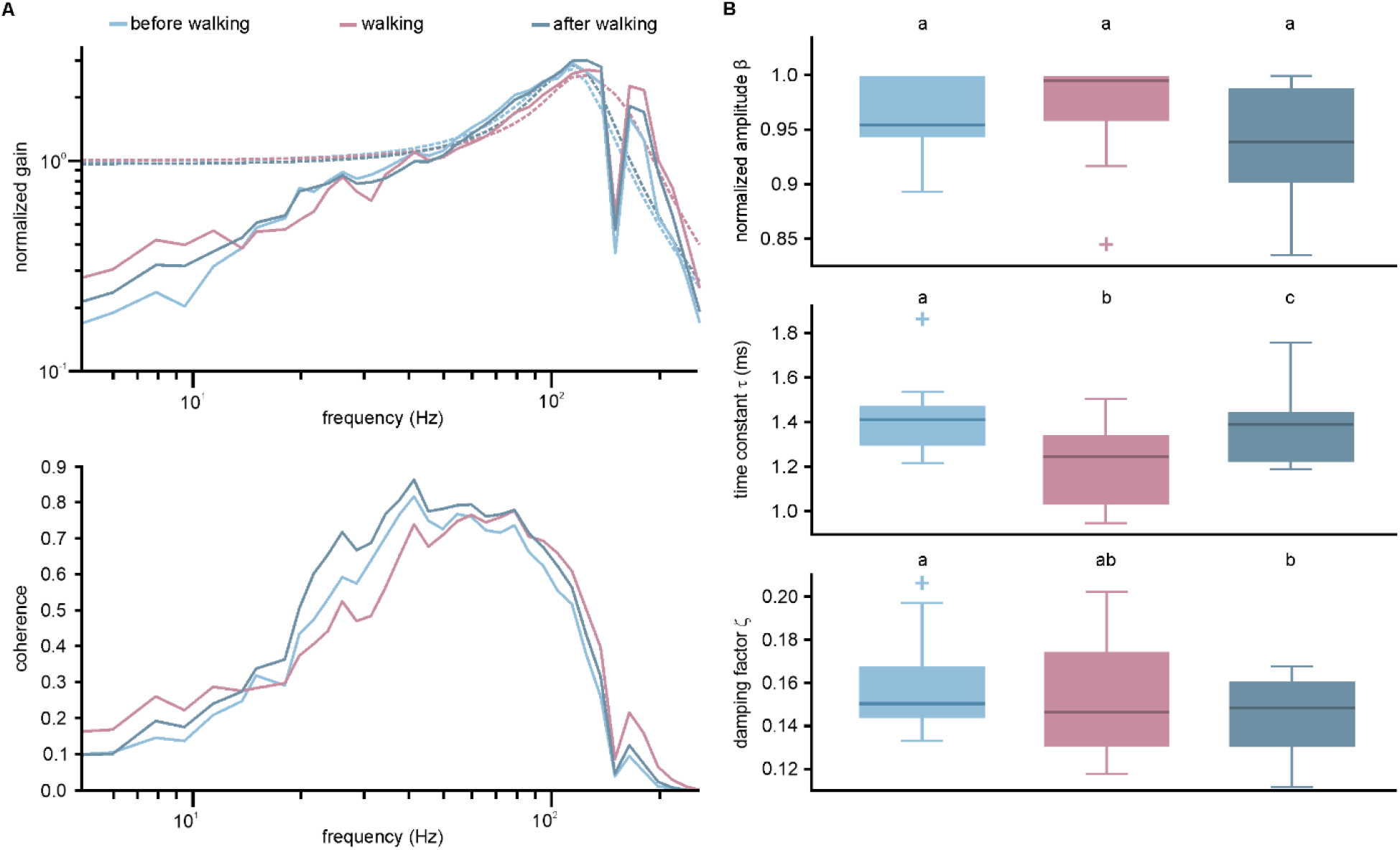
Gain and linear coherence between the stimulus and the ERG for walking experiments. A) To assess which frequency range of the ERG response was specifically affected we calculated gain (upper panel) and linear coherence (lower panel) and averaged over 11 animals. The gain was fitted with a second order low-pass filter (dashed lines). During walking, a shift to higher frequencies was observed. B) Parameters of second-order low-pass fit to the gain. The amplitude β did not differ between any of the conditions (upper panel; p > 0.05, Wilcoxon-Test with Bonferroni correction). The time constant τ became significantly smaller during walking (middle panel; p = 0.001 Wilcoxon-Test with Bonferroni correction). There was a small, but significant difference in τ between the two sitting conditions (middle panel; p = 0.014 Wilcoxon-Test with Bonferroni correction). Walking had no effect on the damping factor ζ (lower panel; p > 0.1, Wilcoxon-Test with Bonferroni correction), but there was a small, but significant difference between the two sitting conditions (lower panel; p = 0.004, Wilcoxon-Test with Bonferroni correction. Boxplots show median, interquartile range (IQR), whiskers with 1.5x IQR and outliers >1.5x IQR.

To test if the changes in visual processing speed can be explained by changes in temperature, we used a thermographic camera to track the temperature of the bumblebee’s compound eyes while sitting and during walking. We found that during walking the temperature of the animals’ eyes increased in conjunction with the heating of their thorax, and then decreased again when the animals were sitting (Figure 2D, Supplemental Figure 2, Supplemental Video). Comparison of the changes in time lag and temperature revealed that both effects occurred during walking and slowly decreased as soon as the animal stopped walking. These temperature time lag correlations applied to all animals tested (Figure 2C).

### Experimental increase of eye temperature replicates effects of walking

To test whether the changes in temperature of the compound eyes during walking were sufficient to explain the observed changes of the visual processing speed, ERG recordings using Gaussian white noise stimulation were carried out as before but this time, the animals were not walking during the experiment. Instead, their head temperature was experimentally raised up to 37°C using an infrared heating lamp. Again, the cross-correlation between the Gaussian white noise stimulus and the ERG was calculated to determine the time lag between the two signals. The example cross-correlations in Figure 4A illustrate that the time lag in the heated condition was smaller (orange; 7.2 ms) compared to the initial temperature before heating and when returning to that temperature after heating (light and dark blue; both 10.8 ms). The median time lag of all animals was significantly smaller (Figure 4B; n = 22; p = 3*10^−5^, Wilcoxon-Test with Bonferroni correction (α = 0.0167)) when the animals were heated up (31.9°C ± 2.0°C) than when they had their initial temperatures (before: 26.9°C ± 1.2°C; after: 27.0°C ± 1.8°C).

**Figure 4:**
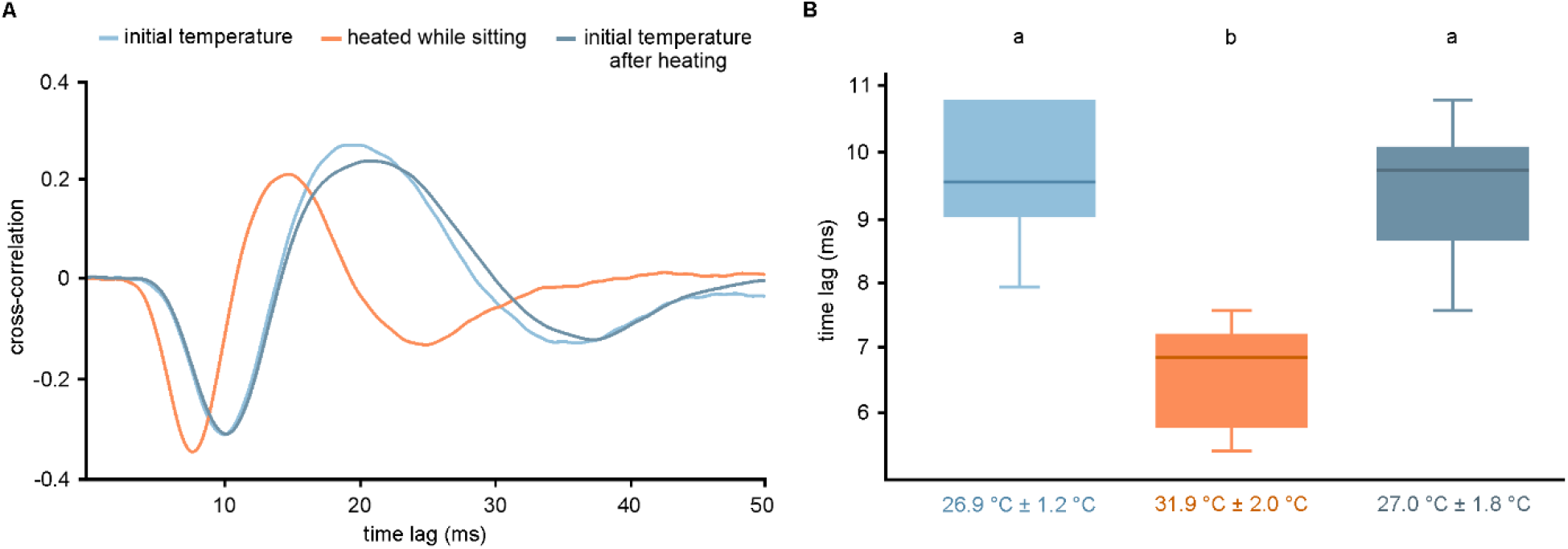
Effects of temperature on electroretinograms of photoreceptor cells in bumblebees stimulated with noise stimulus. (A) Cross-correlation between the signal of the photodiode and the ERG to measure the time lag of the ERG during stimulation with Gaussian noise. The minimum of the cross-correlation occurred earlier, when the bumblebee was heated up while it was sitting (orange) compared to when the animal was just sitting (shades of blue). (B) Boxplots show the responses of 22 animals to the noise track, before heating (light blue; median temperature: 26.9°C ± 1.2°C), while heated (orange; 31.9°C ± 2.0°C), and after return to initial temperature (dark blue; 27.0°C ± 1.8°C). The time lag was significantly shorter when heated than at the initial temperature (p = 3*10^−5^, Wilcoxon-Test with Bonferroni correction), but no differences were seen when comparing the initial temperatures before and after heating experiments (p = 0.05, Wilcoxon-Test with Bonferroni-correction). Boxplots show median, interquartile range (IQR), whiskers with 1.5x IQR and outliers >1.5x IQR.

Furthermore, gain and coherence showed a clear shift to higher frequencies in heated animals (Figure 5A). The time constant τ of the second-order low-pass filter that was fitted to the data was significantly smaller during heating compared to both sitting conditions (Figure 5B middle panel; n = 22; median values: before heating, 1.48 ms, heating, 1.14 ms, after heating, 1.45 ms; before heating vs. heating: p < 0.0001, after heating vs. heating: p < 0.0001, before heating vs. after heating: p = 0.592, Wilcoxon-Test with Bonferroni correction (α = 0.0167)). The damping factor ζ was significantly increased during heating compared to both sitting conditions (initial temperature before and after heating), while it did not differ between the initial temperature conditions. (Figure 5B lower panel; n = 22; median values: before heating, 0.165, heating, 0.184, after heating, 0.148; before heating vs. heating: p = 0.0003, after heating vs. heating: p = 0.0010, before heating vs. after heating: p = 0.287, Wilcoxon-Test with Bonferroni correction (α = 0.0167)). Heating of the animals also increased the amplitude β of the fit function significantly compared to the sitting before heating condition, whereas the amplitude between the sitting conditions and the sitting after heating compared to heating showed no significant difference (Figure 5B upper panel; n = 22; median of normalized values: before heating, 0.897, heating, 1.000, after heating, 0.932; before heating vs. heating: p = 0.0012, after heating vs. heating: p = 0.0309, before heating vs. after heating: p = 0.0261, Wilcoxon-Test with Bonferroni correction (α = 0.0167)).

**Figure 5:**
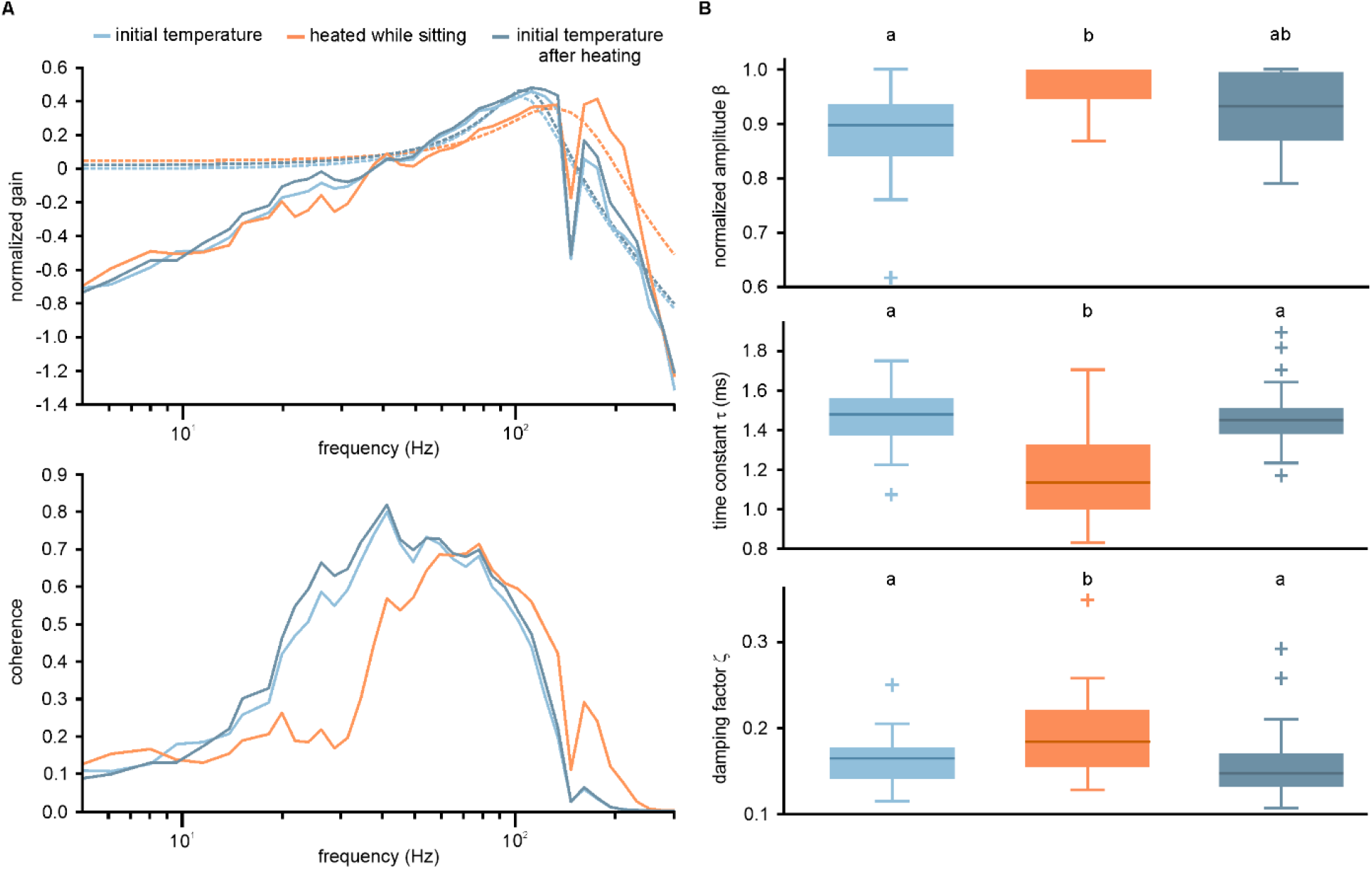
Gain and linear coherence between the stimulus and the ERG for heating experiments. A) Averaged gain and coherence showed a clear shift to higher frequencies in heated animals (n = 22). Dashed lines are fits of second order low pass filter functions. B) Parameters of second order low pass filter fits. The amplitude β of the fit function increased significantly during heating compared to the condition before heating (p = 0.0012, Wilcoxon-Test with Bonferroni correction) whereas the amplitude there were no significant differences between the sitting conditions and between the heating condition and the after heating condition (p > 0.02, Wilcoxon-Test with Bonferroni correction). The time constant τ was significantly smaller during heating compared to both sitting conditions (p < 0.0001, Wilcoxon-Test with Bonferroni correction), but showed no significant difference between the sitting conditions (p = 0.592, Wilcoxon-Test with Bonferroni correction). The damping factor ζ was significantly increased during heating compared to both sitting conditions (p ≤ 0.001, Wilcoxon-Test with Bonferroni correction) while it did not differ between the sitting conditions (p = 0.287, Wilcoxon-Test with Bonferroni correction). Boxplots show median, interquartile range (IQR), whiskers with 1.5x IQR and outliers >1.5x IQR.

To statistically assess if temperature changes are sufficient to explain the increase in visual processing speed in walking bumblebees, we evaluated the time lag between stimulus and response at five different eye temperatures of walking and externally heated bumblebees. For each experiment, we fitted a linear regression to the data points. We then compared the slope of the regression lines between walking (n = 12) and externally heated animals (n = 22) and found no significant difference (p = 0.071, Wilcoxon-Test). We therefore pooled the data from walking and externally heated bumblebees and fitted a linear regression to the entire data set (Figure 6A; y = −0.44x + 21.1; R^2^ = 0.899). It should be noted that, while the linear regression describes the relationship between temperature and processing speed well in our dataset, this is probably only true for a limited temperature range while the effect likely saturates at higher temperatures.

**Figure 6:**
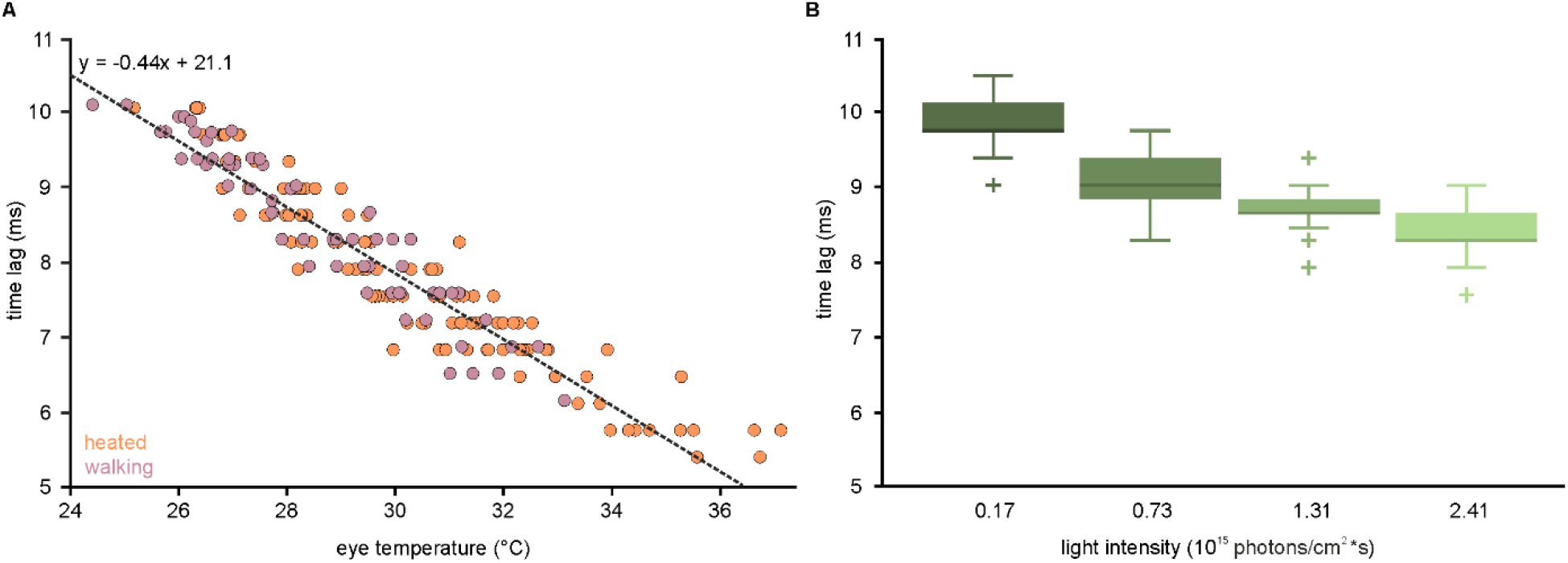
Comparison of temperature and intensity effects. (A) Time lag for walking (violet; 60 data points from 12 animals) and heated (orange; 110 data points from 22 animals) animals were plotted against the eye temperature. With increasing temperature, the time lag of the ERG decreases. The values are well fitted by a linear regression (y = −0.44x + 21.1, R^2^ = 0.899). (B) The time lag (ms) of 17 animals was measured after adaptation to four different intensities (1.74*10^14^ photons/cm^2^*s, 7.27*10^14^ photons/cm^2^*s, 1.31*10^15^ photons/cm^2^*s, 2.41*10^15^ photons/cm^2^*s). As the light intensity increases, the time lag decreases. Boxplots show median, interquartile range (IQR), whiskers with 1.5x IQR and outliers >1.5x IQR.

### Walking accelerates visual responses as much as a 14-fold increase in light intensity

Another factor that can increase the processing speed of photoreceptors is adaptation to higher light intensities (Skorupski and Chittka 2011). We therefore asked how the walking-induced increase in processing speed (from 9.0 ms to 7.4 ms) compares to the increase in processing speed that is induced by adaptation to higher light intensities.

To test this, we presented the Gaussian white noise stimulus to sitting bumblebees at four different light intensities (1.74*10^14^ photons/cm^2^*s, 7.27*10^14^ photons/cm^2^*s, 1.31*10^15^ photons/cm^2^*s, 2.41*10^15^ photons/cm^2^*s). As expected, adaptation to higher light intensities led to a shorter time lag between stimulus and response (Figure 6B). The difference in time lag between the highest and the lowest light intensity tested was 1.4 ms. In comparison locomotion decreased the time lag between stimulus and response by 1.6 ms. In other words, walking increased the processing speed more than a 14-fold increase in light intensity did. This strong effect is especially impressive, when considering that the median observed temperature change during walking was only 2.3°C. It is known that the head temperature of bumblebees in flight can rise up to 35°C (Reber et al. 2015). From our data, we conclude that the time lag between stimulus and ERG in a flying bumblebee would be 5.7 ms, i.e. 3.3 ms faster than during sitting (Figure 6A).

## Discussion

We show here that the ERG response of the bumblebee’s compound eye is accelerated during walking and that this increase in visual processing speed is caused by a rise of the compound eye’s temperature. The observed temperature changes of the eyes are tightly linked to temperature changes in the thorax (Supplemental Figure 2), where the flight muscles are used to generate heat. While the idea that active heat production could be used to increase the temporal resolution of the compound eyes has been suggested more than 20 years ago (Tatler et al. 2000), our study provides the first direct experimental evidence showing that the heat that an insect produces in its thorax during walking is indeed sufficient to effectively speed up the visual system.

Studies in flies have shown that the behavioral state increases the gain of motion-sensitive visual interneurons (Maimon et al. 2010; Chiappe et al. 2010). These changes are mediated through the release of octopamine by modulatory interneurons (Rien et al. 2012; Suver et al. 2012). Recently, it was shown that octopamine signaling is also necessary for shivering thermogenesis in the flight muscles of honeybees (Kaya-Zeeb et al. 2022) and this is likely to also apply to bumblebees. Like the gain increase in visual interneurons, the increase in photoreceptor processing speed is therefore likely to also involve the action of octopamine. While octopamine acts through very different pathways in these two cases, the resulting action is always an adjustment of the visual system to the needs of the current locomotor state.

Neurophysiological processes depend on temperature because electrochemical neuronal potentials are established by asymmetric distribution of ions across the cell membrane. It is therefore not surprising, that sensory processes are temperature dependent. However, the way in which temperature influences sensory processes is modality-dependent. For example, in mechanoreceptors of cockroaches and spiders, higher temperatures lead to an increase in response gain that is independent of stimulus frequency (French and Kuster 1982; Höger and French 1999). This means that a temperature change, which doubles the gain at a low stimulus frequency, will also double the gain at high frequencies, but the range of frequencies that the receptor responds to is not changed. Photoreceptors, on the other hand, usually become more sensitive to higher frequencies with increasing temperature (Tatler et al. 2000; French and Järvilehto 1978; Juusola and Hardie 2001). Higher temperatures of a receptor can be either caused by a rise in ambient temperature or through production of heat by the insect. As locomotion speed of insects is usually positively correlated with ambient temperature, both cases are usually linked to a higher locomotion speed. While locomotion inevitably increases the range of temporal frequencies that the eyes have to process, it is unlikely to have a strong effect on the frequency range, which mechanoreceptors might be exposed to. Physiological changes that the two receptor systems undergo during temperature changes, seem therefore be matched to the behavioral needs of the animals. It is therefore conceivable that walking (and flying) insects actively increase the temperature of their eyes in order to adjust the frequency range of their receptors to the sensory needs. Visually guided flight requires an even faster response of the compound eyes than walking. Reber et al. (2015) found head temperatures of up to 35°C in flying bumblebees, suggesting even faster visual responses than the ones we measured in walking animals. While the activity of flight muscles always leads to heat production in the thorax, it should be mentioned that the blood flow to the head, and therefore its temperature can be actively regulated (Heinrich 1993).

Active increase of eye (and brain) temperature has been previously shown in large pelagic fish, like marlins, sailfish, and sawfish. These species can swim in cold water and have developed a special tissue to heat up their eyes about 10°C above the ambient temperature (Block 1986, Carey 1982). In the swordfish retina, such a rise in temperature effectively increases the processing speed of visual stimuli in the retina by a factor of 5.2 when the eye is heated up by 10 degrees (Fritsches et al. 2005). It is assumed that active heating of the eyes gives these predatory fish an advantage over their prey, when hunting in cooler waters. Similar to the mechanism in sawfish, heat production of bumblebees during walking could be an active way of tuning their eyes to the sensory requirements of the current behavior.

Faster processing of sensory information at the receptor level will only be useful to the animal if the subsequent neuronal circuitry is also adapted to handle the faster incoming information. While our observations in bumblebees pertain only to the primary sensory processing of visual information, heating of the head will necessarily increase the temperature of the brain as well and thus effect any aspect of neuronal processing. It is well documented through electrophysiological recordings from visual interneurons in insects, that a rise in temperature leads to a decrease in response latency (Warzecha et al. 1999, Simmons 2011) and an increase in firing rate (Spavieri et al. 2010). Of course, such effects are not limited to the visual system as temperature effects basic neuronal properties like the input resistance, the membrane time constant and hence the conduction velocity, as well as synaptic transmission (Burrows 1998, Xu and Robertson 1994). The overall effect of temperature on complex neuronal circuits is hardly predictable. For example, central pattern generators (CPGs) in the same animal can respond very differently to changes in temperature. While the locust flight CPG is almost completely temperature-compensated, the locust ventilation CPG increases its frequency 2.3 fold when temperature rises by 10°C (Robertson and Money 2012). It would therefore be desirable for future neuroethological electrophysiology experiments to take the natural temperature range that is linked to the behavior under investigation into account, whenever possible.

When we compared the effect of temperature and adaptation on the increase in response speed, we found that that walking had a similar effect as a 14-fold increase in light intensity. While our experiments were done with rather high light intensities on light-adapted animals, bumblebees are known to be able to also fly under dim light conditions (Reber et al. 2015). However, light levels seem to be a limiting factor for the flight velocity of the bumblebees. The darker the ambient light levels, the slower the velocity of the animals and the more tortuous their flight paths (Reber et al. 2015). The ability to fly at all at low light levels might be a result of the ability to heat, and hence speed up the visual system, to compensate for a reduction in processing speed that is necessitated by longer integration times in dim light.

Our findings that heat produced during walking accelerates the visual system of bumblebees is very likely to be also of profound significance to many other insect species that are capable of temperature regulation (Heinrich 1993). It will be therefore interesting to see how increased head temperatures during walking or flying influence the visual system and the neuronal responses of other insects like large moths, beetles, or dragonflies.

## Material and Methods

### Animals

All experiments were performed using adult, worker bumblebees (*Bombus terrestris*). Colonies were obtained from a commercial supplier (Biobest Group NV, Westerlo, Belgium) and kept either in climate chambers at 25 °C and 55% rH under 12:12 LD or in flight arenas in a laboratory room with windows. Animals were provided with *ad-libitum* food (pollen (Naturwaren-Niederrhein GmbH, Goch, Germany) and API-invert (Biobest Group NV, Westerlo, Belgium)).

Prior to electrophysiological recordings, bumblebees were placed in the freezer for 5–20 min to immobilize them. The animals were then waxed to a 3D-printed holder using dental wax (Omnident, Rodgau, Germany). Head and anterior edge of the thorax were waxed to the holder to prevent head movement-induced recording artifacts, while legs and abdomen were free to allow walking.

ERGs were recorded differentially, using two silver wires (Ø 0.075 mm, Advent Research Materials Ltd, Eynsham, UK) as recording electrodes that were each inserted into one of the compound eyes as superficially as possible. Prior to insertion, the cornea of each compound eye was punctured with a minute pin (Ø 0.2 mm, Ento Sphinx s.r.o., Pardubice, Czech Republic). The insertion site was afterwards sealed with petroleum jelly to protect the eye from desiccation and to stabilize the electrodes mechanically. At the end of the preparation the reference electrode (silver wire, Ø 0.25 mm, World Precision Instruments, Sarasota, USA) was inserted into the head capsule at the posterior dorsal edge of the head just outside the compound eye.

### Electroretinograms

All experiments were carried out at ambient temperatures of 25°C. The extracellular signals were highpass filtered at 1 Hz and amplified 100x with an ELC-01MX amplifier (npi electronic, Tamm, Germany), digitized with a Power 1401 (Cambridge Electronic Design, Cambridge, UK) at 3 kHz, and recorded with Spike2 version 9 (Cambridge Electronic Design, Cambridge, UK).

### Behavioral Setup

Tethered bumblebees were held by a micro manipulator and positioned to sit in a natural posture on an air supported Styrofoam ball (diameter: 50 mm) in the setup. The forward motion and rotation in the yaw axis of the ball were registered using an optical mouse sensor connected to an Arduino Due and transferred to the CED 1401 as an analog signal. The temperature of the animals’ eye was continuously monitored using a thermographic camera (FLIR A65, lens: 45°, *f* = 13 mm, FLIR, Wilsonville, USA). The thermographic data were recorded, stored and extracted using FLIR software. A schematic view of the complete experimental setup is shown in Fig. 1A.

### Stimulation

For visual stimulation light from a green LED (Osram Oslon SSL 80 LT CP7P, dominant wavelength: 528 nm, full width at half maximum: 33 nm) was focused onto one end of a light guide. The other end was positioned 9 cm away from the eye of the animal. The current through the LED was controlled by a custom-built voltage-to-current-converter which was driven by an analog output of the CED 1401. Stimulus intensity was calibrated using a spectrophotometer (Maya 2000 Pro, Ocean Optics). During experiments, light stimuli were monitored by recording the output from an integrated photodiode/transimpedance amplifier, which was placed across from the light guide. In the first set of experiments animals were stimulated with light pulses (50 ms duration at 1 Hz, pulse intensity: 2*10^15^ photons/cm^2^*s). In the second set of experiments, we used Gaussian white noise as a stimulus. A 10 s Gaussian White noise signal was created using the open-source software Audacity (www.audacityteam.org) and saved as a .wav file, which was then imported into Spike2. The noise was then low-pass filtered using the Spike2 implementation of a 9-pole Butterworth filter with a corner frequency of 250 Hz. This gave a noise stimulus with a flat power spectrum up to approximately 220 Hz (see Supplemental Figure 1). Absolute intensity of the stimulus was adjusted using neutral density filters. In most recordings the mean intensity of the noise stimulus was set to 1.74*10^14^ photons/cm^2^*s. To test the intensity dependence of the ERG response three additional mean intensities were used: 7.27*10^14^ photons/cm^2^*s, 1.31*10^15^ photons/cm^2^*s, 2.41*10^15^ photons/cm^2^*s. Prior to testing, animals were adapted for at least ten minutes with the intensity that was to be used during testing. To compare different states of locomotion, the animals first sat on the Styrofoam ball without walking, then were encouraged to walk by gentle brush strokes on the abdomen (some animals started walking without encouragement) and at the end sat again. Visual stimulation was continuous throughout each experiment.

To test the effect of externally applied heat the infrared heating lamp was turned on until the animals’ eye had a temperature of about 37°C. Then the heat lamp was turned off and recording was continued until the initial head temperature was reached.

### Quantification and statistical analysis

All experiments were evaluated using custom-written Spike2 scripts.

### Pulses

Responses to light pulses were evaluated by extracting the times of the light pulses and the times when the ERG amplitude crossed a threshold of −2.5 mV. The difference between the onset of each light pulse and the crossing of the threshold gave the time lag. Time lags were averaged for 5 seconds (i.e. from 5 pulses) either from the phase immediately prior to the onset of walking, during walking or after a recovery period of at least 5 minutes.

### Cross-correlation

To measure the time lag of the ERG during stimulation with Gaussian noise we calculated the cross-correlation between the signal of the photodiode and the ERG. Cross-correlation measures the similarity of two waveforms in the time domain and can assume values between −1 and 1. A value of 1 indicates identical waveforms (except for amplitude), 0 indicates no correlation between the waveforms and −1 indicates identical waveforms with inverted signs (except for amplitude). The x-value at the minimum of the cross-correlation indicates the time lag between the two curves.

During experiments, cross-correlations were continuously calculated to monitor the ERG time lag. For evaluation, we chose segments of 5 s immediately preceding walking (or application of an external heat stimulus), immediately after walking (or heat stimulus) and upon recovery after at least 5 minutes. Walking activity often caused artifacts in the ERG recording, which made it necessary to choose a post-walking segment for evaluation, rather than the last 5 s of walking activity. Because of the slow time-course of recovery, this had only minor influence on the results.

### Coherence

Linear coherence was used to measure the similarity in frequency content between the white noise light stimulus measured by the photodiode and the ERG. At any given frequency, coherence is 1 if the relationship between input and output is purely linear, i.e. if input and output waveform have a constant amplitude ratio and phase over the evaluated time period and the system is free of noise. Values smaller than 1 indicate either nonlinearity or added noise. The linear coherence function γ^2^(f) ranges between 0 and 1. It was calculated as:

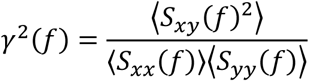

where S_xx_(f) is the input spectrum (of Gaussian white noise stimulus measured by the photodiode), S_yy_(f) is the output spectrum (of ERG), S_xy_(f) is the cross-spectrum, and 〈〉 indicates averaging over ensembles (Bendat and Piersol 1980). Spectra were obtained by resampling the data at 1 ms sampling intervals and calculating the Fast-Fourier-Transformation (Cooley and Tukey 1965) using 512 point segments.

### Gain and fitting

Frequency responses were obtained by direct spectral estimation from the cross-spectrum and the input spectrum (Bendat and Piersol 1980) as:

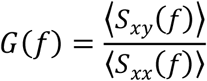

where S_xx_(f) is the input spectrum (of the Gaussian white noise stimulus as measured by the photodiode), S_xy_(f) is the cross-spectrum, and 〈〉 indicates averaging over ensembles. The gain was fitted with a second order low-pass filter of the form:

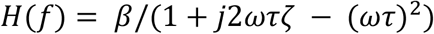

where β is amplitude, τ is a time constant, ζ is a damping factor, ω is radial frequency (2πf) and *j* is √(−1).

### Statistics

All statistical evaluations were done using Matlab (The Mathworks, Natick, MA, USA).

## Acknowledgments

We are grateful to Dr. Andrew French for the evaluation of frequency response data. We thank Dr. Flavio Roces for insightful discussions and for providing the infrared heating lamp, Dr. Randolf Menzel for providing the micro manipulator and the silver wire used in this study, and Dr. Basil el Jundi for helpful comments on an earlier version of the manuscript. This work was supported by a grant from the German Research Council (DFG: PF714/5-1) to K.P.

## Author Contribution

Conceptualization, L.R. and K.P.; Methodology, K.P., L.R., and R.M.; Software, K.P.; Formal Analysis, R.M., E.K., and L.R.; Validation, L.R. and K.P.; Investigation, R.M. and E.K.; Resources, S.K.-Z. and M.T.; Data Curation, L.R. and K.P., Writing – Original Draft, L.R., R.M., and K.P.; Writing – Review and Editing, all authors; Visualization, L.R. and R.M.; Supervision, K.P. and L.R.; Project Administration, K.P.; Funding Acquisition, K.P.

## Declaration of interests

The authors declare no competing interests.

## Notes

### Competing Interest Statement

The authors have declared no competing interest.

## References

Bendat JS, Piersol AG (1980) Engineering applications of correlation and spectral analysis. New York: Wiley: 302 p

Block BA (1986) Structure of the brain and eye heater tissue in marlins, sailfish and spearfish. J Morphol 190:169–189. https://doi.org/10.1002/jmor.1051900203

Burrows M (1989) Effect of temperature on a central synapse between identified motor neurons in the locust. J Comp Physiol A 165:687–695. https://doi.org/10.1007/BF00611000

Cano M, Bezdudnaya T, Swadlow HA, Alonso JM (2006) Brain state and contrast sensitivity in the awake visual thalamus. Nat Neurosci 9:1240–1242. https://doi.org/10.1038/nn1760

Cao L, Händel B (2019) Walking enhances peripheral visual processing in humans. PLoS Biol 17:1–23. https://doi.org/10.1371/journal.pbio.3000511

Carey FG (1982) A brain heater in the swordfish. Science 216:1327–1329. https://doi.org/10.1126/science.7079766

Cheng KY, Frye MA (2020) Neuromodulation of insect motion vision. J Comp Physiol A Neuroethol Sensory Neural Behav Physiol 206:125–137. https://doi.org/10.1007/s00359-019-01383-9

Chiappe ME, Seelig JD, Reiser MB, Jayaraman V (2010) Walking modulates speed sensitivity in drosophila motion vision. Curr Biol 20:1470–1475. https://doi.org/10.1016/j.cub.2010.06.072

Christensen AJ, Pillow JW (2022) Reduced neural activity but improved coding in rodent higher-order visual cortex during locomotion. Nat Commun 13:1–8. https://doi.org/10.1038/s41467-022-29200-z

French AS, Järvilehto M (1978) The dynamic behavior of photoreceptor cells in the fly in response to random (white noise) stimulation at a range of temperatures. J Physiol 274:311–322. https://doi.org/10.1113/jphysiol.1978.sp012149

French AS, Kuster, JE (1982) The effects of temperature on mechanotransduction in the cockroach tactile spine. J Comp Physiol 147:251–258. https://doi.org/10.1007/BF00609849

Fritsches KA, Brill RW, Warrant EJ (2005) Warm Eyes Provide Superior Vision in Swordfishes. Curr Biol. 15:55–58. https://doi.org/10.1016/j.cub.2004.12.064

Heinrich B (1993) The Hot-Blooded Insects: Strategies and Mechanisms of Thermoregulation. Cambridge, MA and London, England: Harvard University Press https://doi.org/10.4159/harvard.9780674418516

Höger U, French AS (1999) Temperature sensitivity of transduction and action potential conduction in a spider mechanoreceptor. Pflügers Arch Eur J Physiol 438:837–842. https://doi.org/10.1007/s004249900113

Juusola M, Hardie RC (2001) Light adaptation in Drosophila photoreceptors: II. Rising temperature increases the bandwidth of reliable signaling. J Gen Physiol 117:27–41. https://doi.org/10.1085/jgp.117.1.27

Kaya-Zeeb S, Engelmayer L, Straßburger M, Bayer J, Bähre H, Seifert R, Scherf-Clavel O, Thamm M (2022) Octopamine drives honeybee thermogenesis. eLife 11:1–22. https://doi.org/10.7554/eLife.74334

Kikuchi R, Naito K, Minagawa S (1961) Effect of Temperature on the Retinal Slow Potential of the Horseshoe Crab. Nature 190:1011–1012. https://doi.org/10.1038/1901011a0

Maimon G, Straw AD, Dickinson MH (2010) Active flight increases the gain of visual motion processing in Drosophila. Nat Neurosci 13:393–399. https://doi.org/10.1038/nn.2492

Maimon G (2011) Modulation of visual physiology by behavioral state in monkeys, mice, and flies. Curr Opin Neurobiol 21:559–564. https://doi.org/10.1016/j.conb.2011.05.001

Murakami M, Kashiwadani H, Kirino Y, Mori K (2005) State-dependent sensory gating in olfactory cortex. Neuron 46:285–296. https://doi.org/10.1016/j.neuron.2005.02.025

Niell CM, Stryker MP (2010) Modulation of visual responses by behavioral state in mouse visual cortex. Neuron 65:472–479. https://doi.org/10.1016/j.neuron.2010.01.033

Polack PO, Friedman J, Golshani P (2013) Cellular mechanisms of brain state-dependent gain modulation in visual cortex. Nat Neurosci 16:1331–1339. https://doi.org/10.1038/nn.3464

Reber T, Vähäkainu A, Baird E, Weckström M, Warrant E, Dacke M (2015) Effect of light intensity on flight control and temporal properties of photoreceptors in bumblebees. J Exp Biol 218:1339–1346. https://doi.org/10.1242/jeb.113886

Rien D, Kern R, Kurtz R (2012) Octopaminergic modulation of contrast gain adaptation in fly visual motion-sensitive neurons. Eur J Neurosci 36:3030–3039. https://doi.org/10.1111/j.1460-9568.2012.08216.x

Rind FC, Santer RD, Wright GA (2008) Arousal facilitates collision avoidance mediated by a looming sensitive visual neuron in a flying locust. J Neurophysiol 100:670–680. https://doi.org/10.1152/jn.01055.2007

Robertson RM, Money TGA (2012) Temperature and neuronal circuit function: compensation, tuning and tolerance. Curr Opin Neurobiol 22:724–734. https://doi.org/10.1016/j.conb.2012.01.008

Roebroek JGH, van Tjonger M, Stavenga DG (1990) Temperature dependence of receptor potential and noise in fly (*Calliphora Erythrocephala*) photoreceptor cells. J Insect Physol 36:499–505. https://doi.org/10.1016/0022-1910(90)90101-K

Rosner R, Egelhaaf M, Warzecha AK (2010) Behavioural state affects motion-sensitive neurones in the fly visual system. J Exp Biol 213:331–338. https://doi.org/10.1242/jeb.035386

Simmons PJ (2011) The effects of temperature on signalling in ocellar neurons of the desert locust, Schistocerca gregaria. J Comp Physiol A Neuroethol Sens Neural Behav Physiol 197:1083–1096. https://doi.org/10.1007/s00359-011-0669-y

Skorupski P, Chittka L (2011) Photoreceptor processing speed and input resistance changes during light adaptation correlate with spectral class in the bumblebee, Bombus impatiens. PLoS One 6: e25989. https://doi.org/10.1371/journal.pone.0025989

Spavieri DL, Eichner H, Borst A (2010) Coding efficiency of fly motion processing is set by firing rate, not firing precision. PLoS Comput. Biol 6: e1000860. https://doi.org/10.1371/journal.pcbi.1000860

Stern M (2009) The PM1 neurons, movement sensitive centrifugal visual brain neurons in the locust: anatomy, physiology, and modulation by identified octopaminergic neurons. J Comp Physiol A 195:123–137. https://doi.org/10.1007/s00359-008-0392-5

Suver MP, Mamiya A, Dickinson MH (2012) Octopamine Neurons Mediate Flight-Induced Modulation of Visual Processing in *Drosophila*. Curr Biol 22:2294–2302. https://doi.org/10.1016/j.cub.2012.10.034

Tatler B, O’Carroll DC, Laughlin SB (2000) Temperature and the temporal resolving power of fly photoreceptors. J Comp Physiol A 186:399–407. https://doi.org/10.1007/s003590050439

Warzecha A, Horstmann W and Egelhaaf M (1999). Temperature-dependence of neuronal performance in the motion pathway of the blowfly calliphora erythrocephala. J Exp Biol 202:3161–3170. https://doi.org/10.1242/jeb.202.22.3161

Weckström M, Järvilehto M, Kouvalainen E, Järvilehto P (1985) Fly photoreceptors and temperature: Relative UV-sensitivity is increased by cooling. Eur Biophys J 12:173–179. https://doi.org/10.1007/BF00254076

Xu H, Robertson RM (1994) Effects of temperature on properties of flight neurons in the locust. J Comp Physiol A 175:193–202. https://doi.org/10.1007/BF00215115

