## Supplemental Data for "Walking bumblebees see faster"

#### Supplementary Data

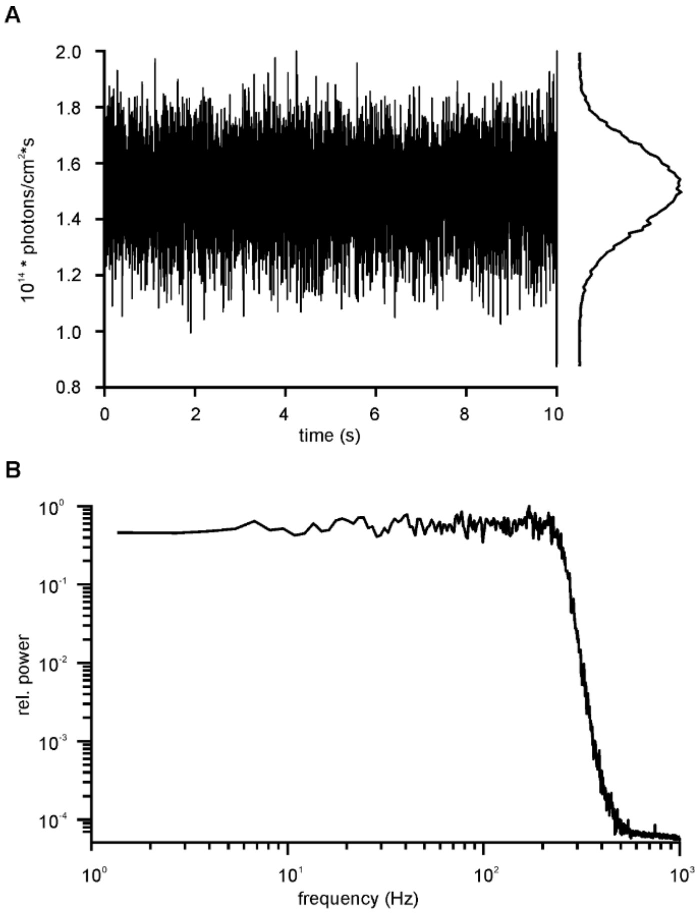


**Figure S1: Properties of the Gaussian white noise stimulus.** A) Whole trace of the 10 s Gaussian white noise stimulus. B) Power spectrum of the Gaussian white noise stimulus.

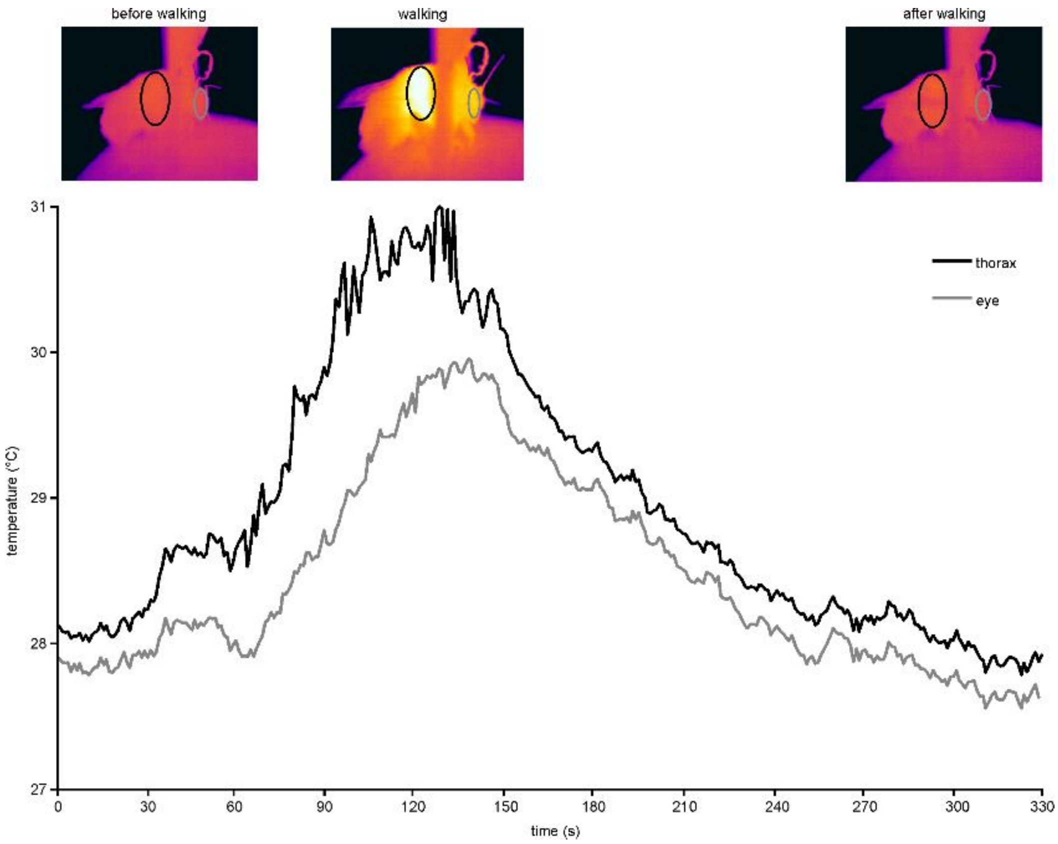


**Figure S2: Thermographic tracking of the Bumblebee.** Top: Thermographic images of a bumblebee before, during and after walking. Gray and black ellipses indicate regions of interest that were used to measure the temperature of the eye and the thorax respectively. Bottom: Time course of temperature measured at the eye (gray) and the thorax (black). During locomotion, both temperatures increased and after walking, the values return to their initial level.

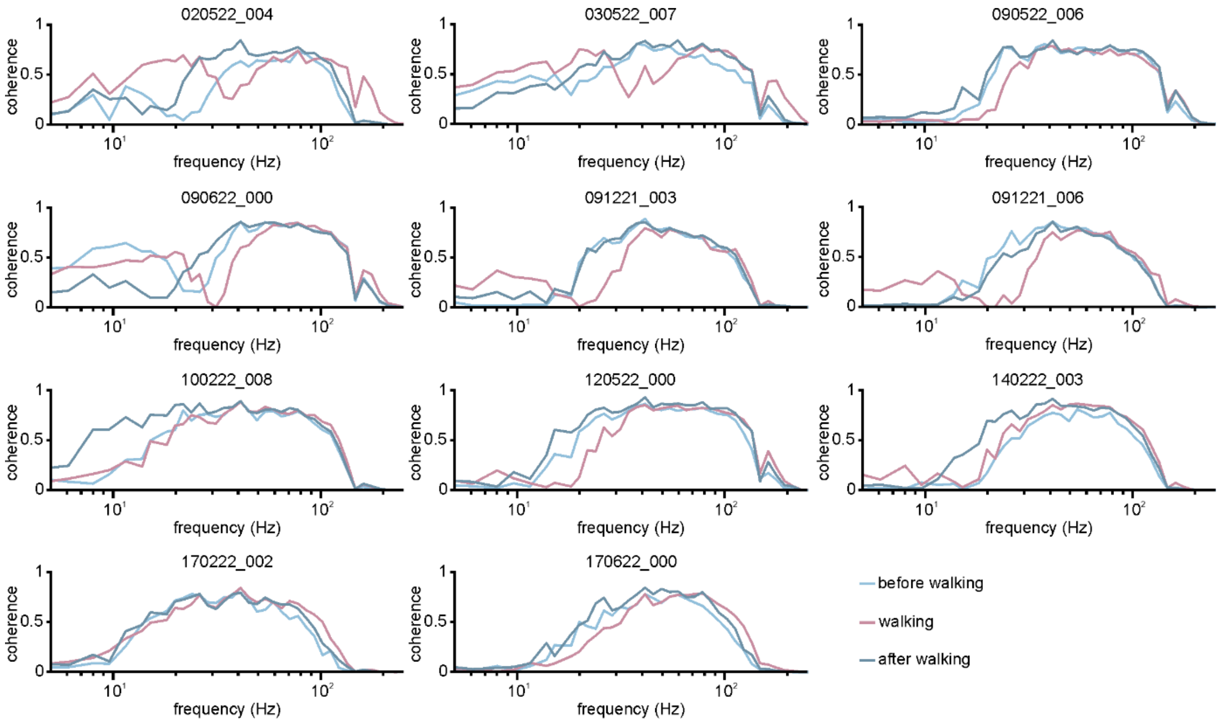


**Figure S3: Coherence between the stimulus and the ERG before, during and after walking.** To assess which frequencies in the ERG response were affected by the changes the linear coherence (n = 11) between the Gaussian white noise stimulus and the ERG was calculated. During walking, coherence was shifted to higher frequencies in most animals.

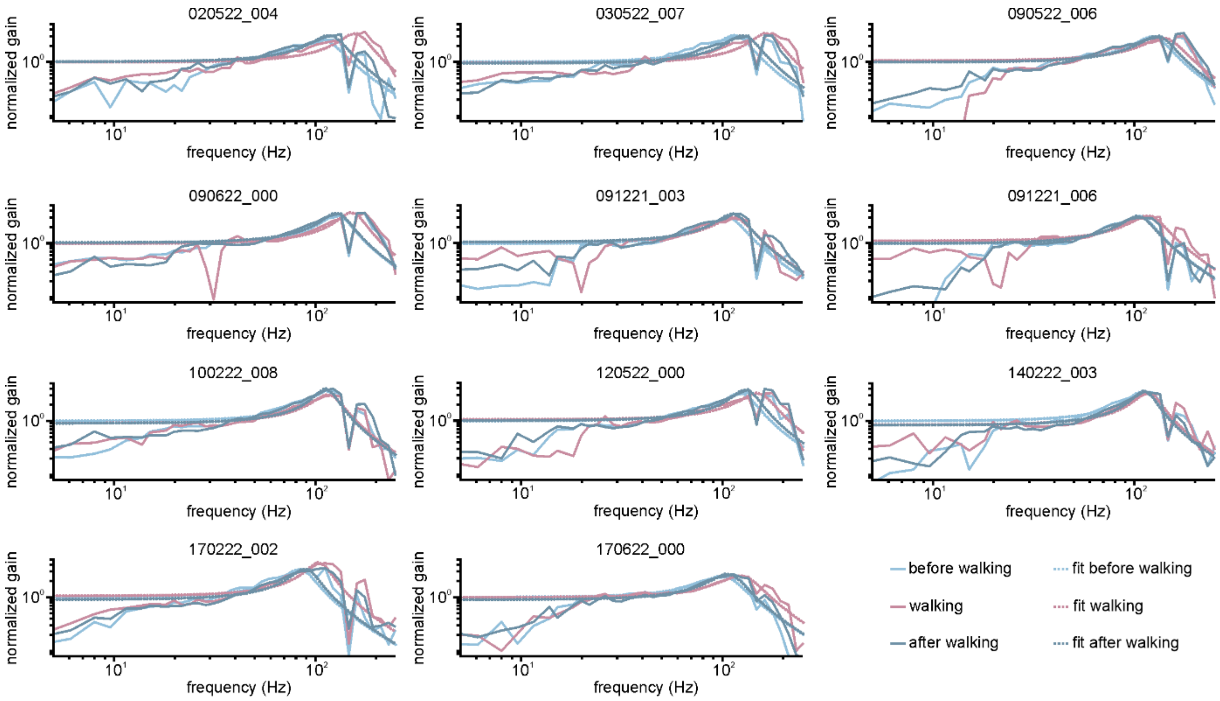


**Figure S4: Gain between the stimulus and the ERG before, during and after walking.** To assess which frequency range of the ERG response was specifically affected we calculated the gain (solid lines) for 11 animals. The gain was fitted with a second order low-pass filter (dashed lines, see Methods). During walking, the roll-off frequency was shifted to higher frequencies in most animals.

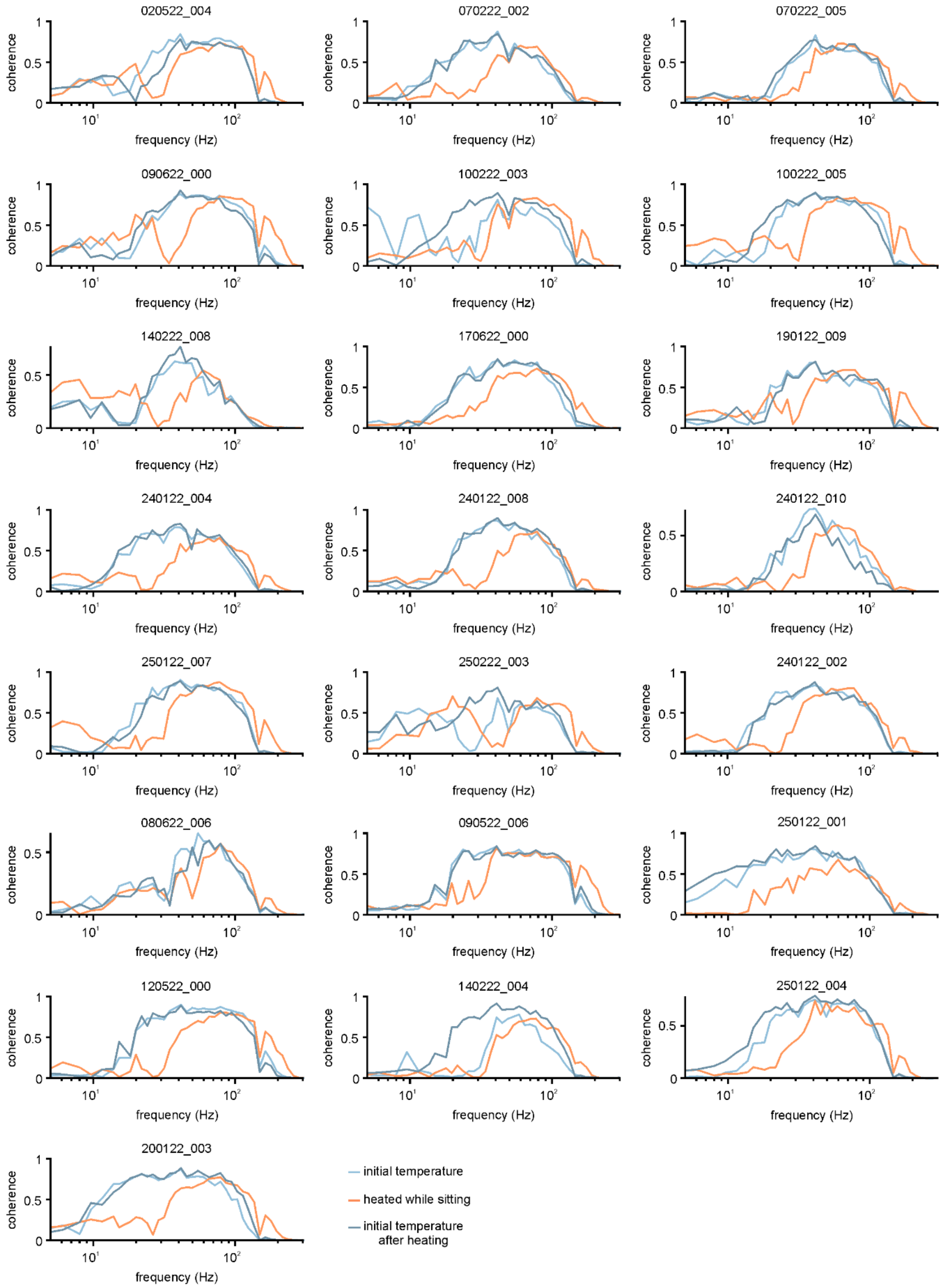


**Figure S5: Coherence between the stimulus and the ERG before, during and after heating.** To assess which frequencies in the ERG response were affected by the changes the linear coherence (n = 22) of the system was calculated. During heating, a shift to higher frequencies was observed.

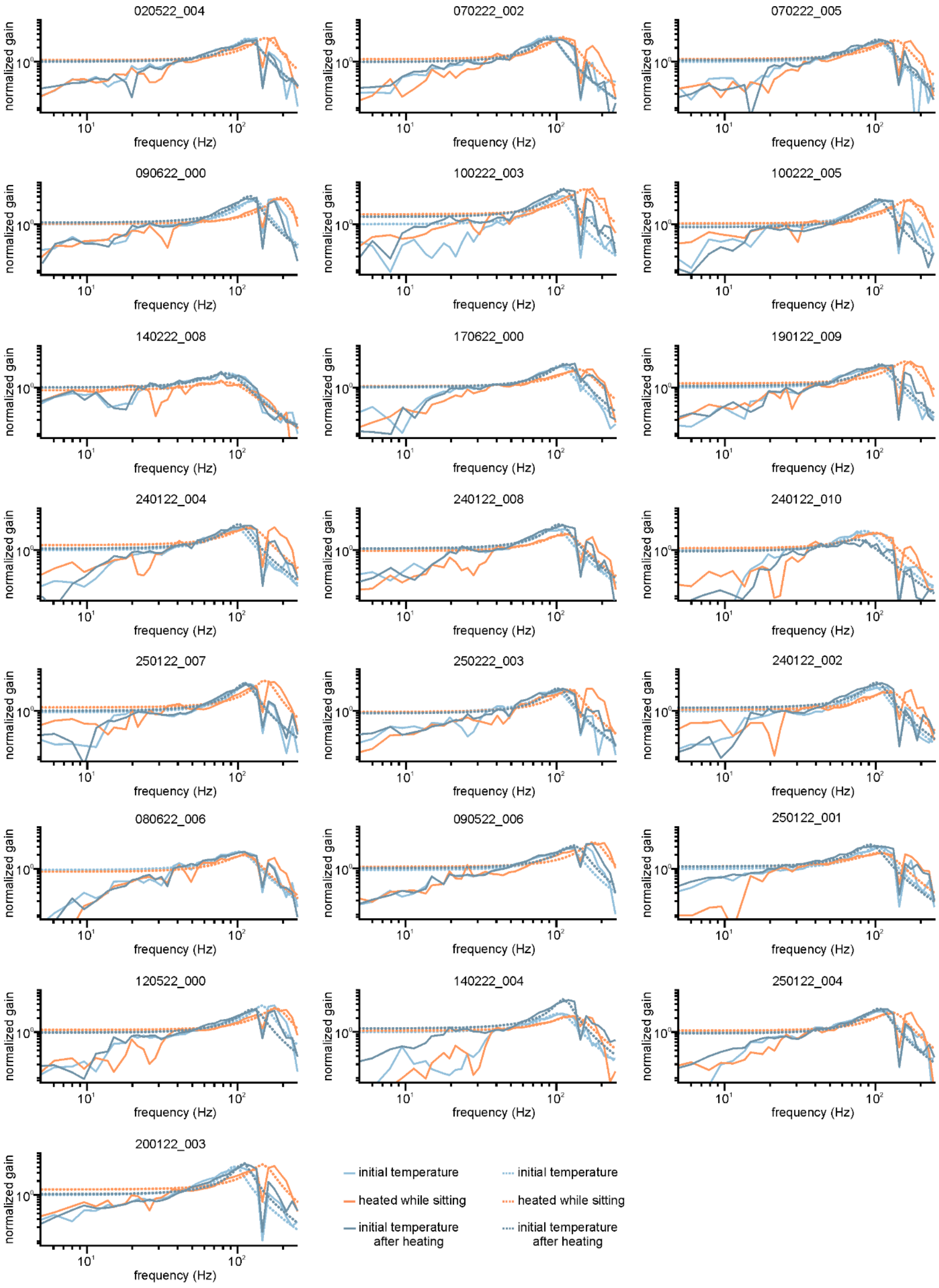


**Figure S6: Gain between the stimulus and the ERG before, during and after heating.** To assess which frequency range of the ERG response was specifically affected we calculated the gain (solid lines) for 22 animals. The gain was fitted with a second order low-pass filter (dashed lines, see Methods). During walking, the roll-off frequency was shifted to higher frequencies.

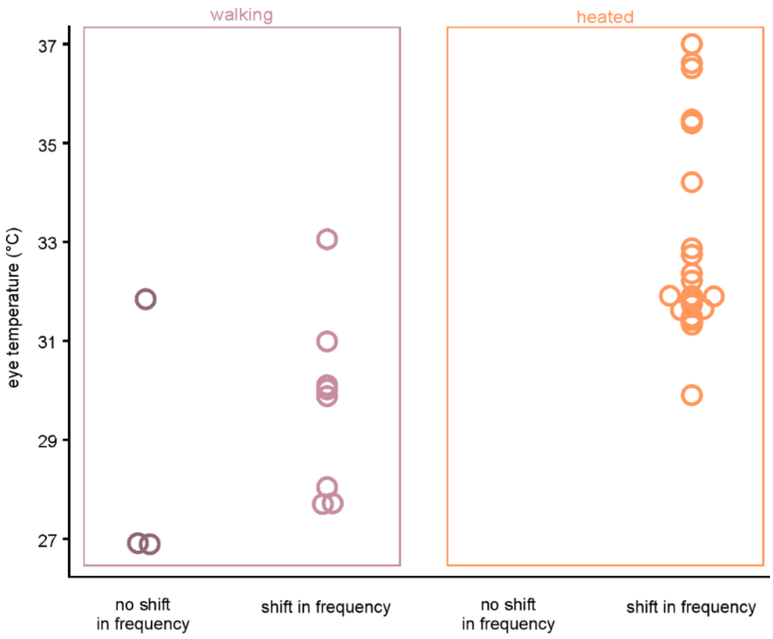


**Figure S7: Comparison of frequency shift of the coherence function with the eye temperature.** A shift in the coherence function was observed in all animals that were heated up by the IR-lamp. In animals that walked, a shift of the coherence function towards higher frequencies was more likely to be observed in animals that heated up more.

### Supplementary Video

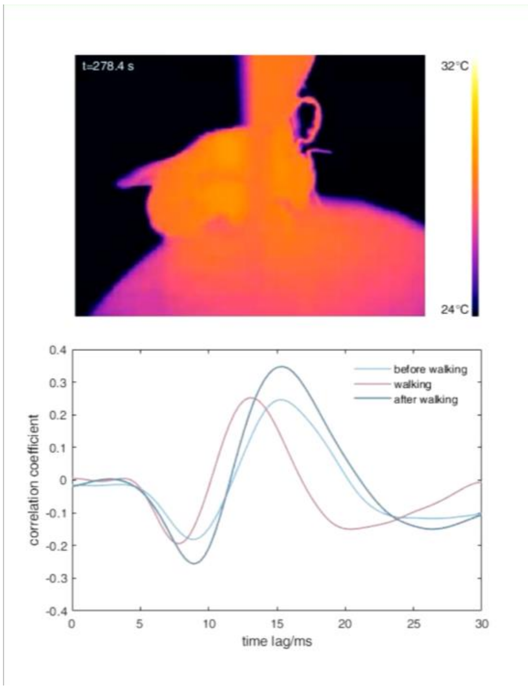


**Supplemental Video 1:** Top: Thermographic video of a tethered bumblebee on an air supported Styrofoam ball. Bottom: Continuous representation of cross-correlation between Gaussian white noise stimulus and electroretinogram. During walking, the temperature of the bumblebees' thorax and head increase and the cross-correlation curve is shifted to the left, indicating a faster response of the visual system. 10x time-lapse.
